# Between Two Extremes: *Tripsacum dactyloides* Root Anatomical Responses to Drought and Waterlogging

**DOI:** 10.1101/2025.03.17.643764

**Authors:** Joel F. Swift, Desi Thimesch, Lucas Bengfort, Shahzaib Asif, Maggie R. Wagner

## Abstract

**Premise:** Plant roots are the critical interface between plants, soil, and microorganisms, and respond dynamically to changes in water availability. Although anatomical adaptations of roots to water stress (e.g., the formation of root cortical aerenchyma) are well documented, it remains unclear whether these responses manifest along the length of individual roots under both water deficiency and water over-abundance.

**Methods:** We investigated the anatomical responses of *Tripsacum dactyloides* L. to both drought and flood stress at high spatial resolution. Nodal roots were segmented into one-centimeter sections from the tip to the base, allowing us to pinpoint regions of maximal anatomical change.

**Results:** Both stressors increased the proportion of root cortical aerenchyma, but metaxylem responses differed: flooding increased vessel area whereas drought led to smaller vessels, with both showing a lower number of vessels. Drought also significantly increased root hair formation, but only within the first two centimeters. The most pronounced anatomical changes occurred 3-7 cm from the root tip, where cortical cell density declined as aerenchyma expanded.

**Discussion:** These findings highlight spatial variation in root anatomical responses to water stress and provide a framework integrating various other data types where sampling effort is limiting (e.g., microbiome, transcriptome, proteome).

## Introduction

While water is a basic requirement for plants, too much or too little can have drastic consequences. As global climate changes, the frequency and severity of extreme weather events have increased (Dethier *et al*., 2020; Davenport *et al*., 2021; Chiang *et al*., 2021; Balting *et al*., 2021), decreasing plant primary productivity and resulting in billions of dollars of losses to crop species (Rosenzweig *et al*., 2002). Plant roots are the point of interaction between plants and soils, serving as the front line in response to dramatic changes in soil water content. Roots under water stress display stunning amounts of phenotypic plasticity, altering their architecture and undergoing dramatic changes in anatomical structure (Karlova *et al*., 2021). Given the role roots play in the absorption of nutrients and water, anchorage, and interactions with soil microorganisms, it will be paramount to further our understanding of their anatomical response to water stress and how this response alters these roles. An impediment to anatomical analysis at the plant phenomics scale (hundreds of samples or more) is the time required to create anatomical sections; therefore, increased sample throughput can be achieved by targeting particular regions of a root with the greater response to water stress.

Water stress for plant roots can take multiple forms from over-abundance (*i*.*e*., flooding and waterlogging) to deficiency (*i*.*e*., drought). Changes to the porosity of roots, particularly the root cortex, tend to be some of the most noticeable anatomical responses to water stress. Greater porosity in the root cortex is achieved by the formation or enlargement of intracellular spaces in the root cortex termed root cortical aerenchyma. Under flooded conditions and waterlogging, roots quickly enter a hypoxic state due to decreased gas diffusion with soil; root cortical aerenchyma can assist in this critical state by allowing diffusion to occur between the root and shoot system (Jackson & Armstrong, 1999). This diffusion can result in higher O_2_ levels in the root tip and allow a route for the release of stress-induced volatiles (*e*.*g*., ethylene, salicylic acid, ethanol, methyl jasmonate) from the shoot (Colmer, 2003; Voesenek & Bailey-Serres, 2015). The increase in cortex porosity is often accompanied by suberization of the root endo/exodermis, which seals the root in a waxy hydrophobic layer decreasing radial oxygen and water loss (Enstone *et al*., 2002; Chen *et al*., 2022). For instance, root exodermal suberin production in *Solanum* improved stem water content and performance under water-deficit conditions (Cantó-Pastor *et al*., 2024). Under drought conditions, increased porosity of roots is hypothesized to decrease the metabolic cost of soil exploration under drought by decreasing the number of living cells to sustain (Zhu *et al*., 2010; Jaramillo *et al*., 2013).

While anatomical responses to water stress are well-documented, most comparative studies have focused on differences among species rather than variation within species or even within individual plants. It remains unclear whether anatomical responses to water stress manifest uniformly along the length of individual roots. This provided the rationale for designing our study, in which we subjected plants to either drought or flood stress and quantified the changes in root anatomy. Addressing this question is essential for understanding how root function varies spatially under stress conditions and for designing experiments that leverage this information.

The objective of this study was to investigate the anatomical responses of *Tripsacum dactyloides* to flood and drought stress along the length of individual nodal roots. Sampling was conducted at a fine resolution by segmenting roots into up to eight one-centimeter segments (from root tip toward the root base). In this way we sought to provide the resolution needed to pinpoint areas of the root that show the greatest response, which will make for fruitful sampling of the microbiome in the future.

Our results showed both commonalities and differences between the water extremes. We found that both flooding and drought increased the proportion of root cortical aerenchyma. While metaxylem vesels were reduced in number under both stresses, the area of vessels increased under flooding while droughted plants showed smaller vessel areas. Only drought caused a significant increase in root hairs, with this relegated to the first two root segments. The greatest degree of change from the control was observed at 3-7 cm from the root tip. Our results suggest that the optimal part of the root for studying drought/flood induced changes in anatomy is 3-7 cm from the root tip. This region experienced the greatest reduction in cortical cell density as the proportion of aerenchyma increased.

## Methods

### Study System

*Tripsacum dactyloides* L. is a warm season perennial bunchgrass native to temperate North America, occupying diverse habitats from prairies and rocky hillsides to streambanks and mesic woodland edges (De Wet *et al*., 1982). Across its range it is noted for its tolerance to drought, flood, and cold (Ray *et al*., 1998; Clark *et al*., 1998; Yan *et al*., 2019). *Tripsacum dactyloides* constitutively forms schizogenous (*i*.*e*., via cellular separation) root cortical aerenchyma (Ray *et al*., 1998), a trait commonly associated with wetland-adapted species (Evans, 2004). Constitutively formed aerenchyma can provide immediate oxygen flow longitudinally to root tips, while also accelerating the formation of additional induced aerenchyma (Gong *et al*., 2019; Pedersen *et al*., 2021b). Because *T. dactyloides* has a close evolutionary relationship with and can be crossed with *Zea mays* L. (Ross-Ibarra *et al*., 2009; Gault *et al*., 2018), it is a potential source of beneficial traits for introgression into maize (Harlan & Wet, 1977; Ray *et al*., 1999; Yan *et al*., 2020).

### Experimental Design and sampling

Seeds of *Tripsacum dactyloides* (L.) cultivar ‘Pete’ were obtained from Gamagrass Seed Company (GERMTEC II; Falls City, NE, USA). ‘Pete’ was developed at the USDA Manhattan Plant Materials Center primarily for forage and silage. It is a composite of 70 accessions from Oklahoma and Kansas that were allowed to open pollinate for three generations (USDA Natural Resources Conservation Service, 2011). Seeds were surfaced-sterilized by soaking in concentrated hydrogen peroxide (30% v/v) for two hours, followed by rinsing three times with distilled water. We have found this method to improve emergence percentage and decrease rates of fungal contamination. Seeds were planted into two 25x50 cm 98-well seed trays; each well was plugged with a sterilized cotton ball and filled with autoclaved calcined clay (“Pro’s Choice Rapid Dry”; Oil-Dri Corporation, Chicago, IL, USA), with each well receiving two seeds. Seed trays were placed into a growth chamber (Conviron PGR15, Pembina, SD, USA) set to 12-hour light/dark cycle at 27/23° C, ambient humidity, and a light intensity of 648 μmol m^-2^s^-1^. We bottom-watered seed trays daily to maintain a constant moisture level. We transplanted seedlings in two germination cohorts, at 10 and 14 DAP (n = 67 and 12, respectively) to pots (1 pint, ∼473ml) filled with peat moss and perlite based potting soil (BM 6; Berger, Saint-Modeste, QC, CAN). Seedlings were grown in the growth chamber for five additional days prior to relocating to a temperature-controlled greenhouse (23/20 ºC day/night, respectively) at the University of Kansas (Lawrence, KS, USA). Plants were arranged under growth lights (250 W Sun Systems Compact Cool fluorescent lamps) to supplement natural light for 12 hours/day during the greenhouse experiment.

Seedlings were monitored and watered every other day prior to commencement of the stress treatments. Data on shoot height at 28 DAP was used to distribute plants into treatment groups (n = 60, 12 per treatment) such that the mean height of plants within each treatment group was equivalent. Treatments were as follows: 10 days without watering (Drought), 24, 48, and 72 hours of soil waterlogging (Waterlogged 24, 48, 72, respectively), and normal watering conditions (Control; Figure 1). All treatment groups had a small amount of sand placed on top of the soil surface, which was necessary to stabilize soil in the waterlogging treatment but was applied to all treatment groups for consistency. Treatments groups were harvested directly following their stress treatment, with no recovery period. Upon harvesting, the shoot system was removed and dried, roots were washed of soil and a single nodal root (borne from the plant base) was collected for anatomical analysis. This nodal root was cut into up to eight one-cm segments, from the tip towards the root base, which were placed into 1.5 mL tubes filled with 75% ethanol (v/v) and stored at 4 ºC until cross-sectioning (Figure 1).

**Figure 1.**
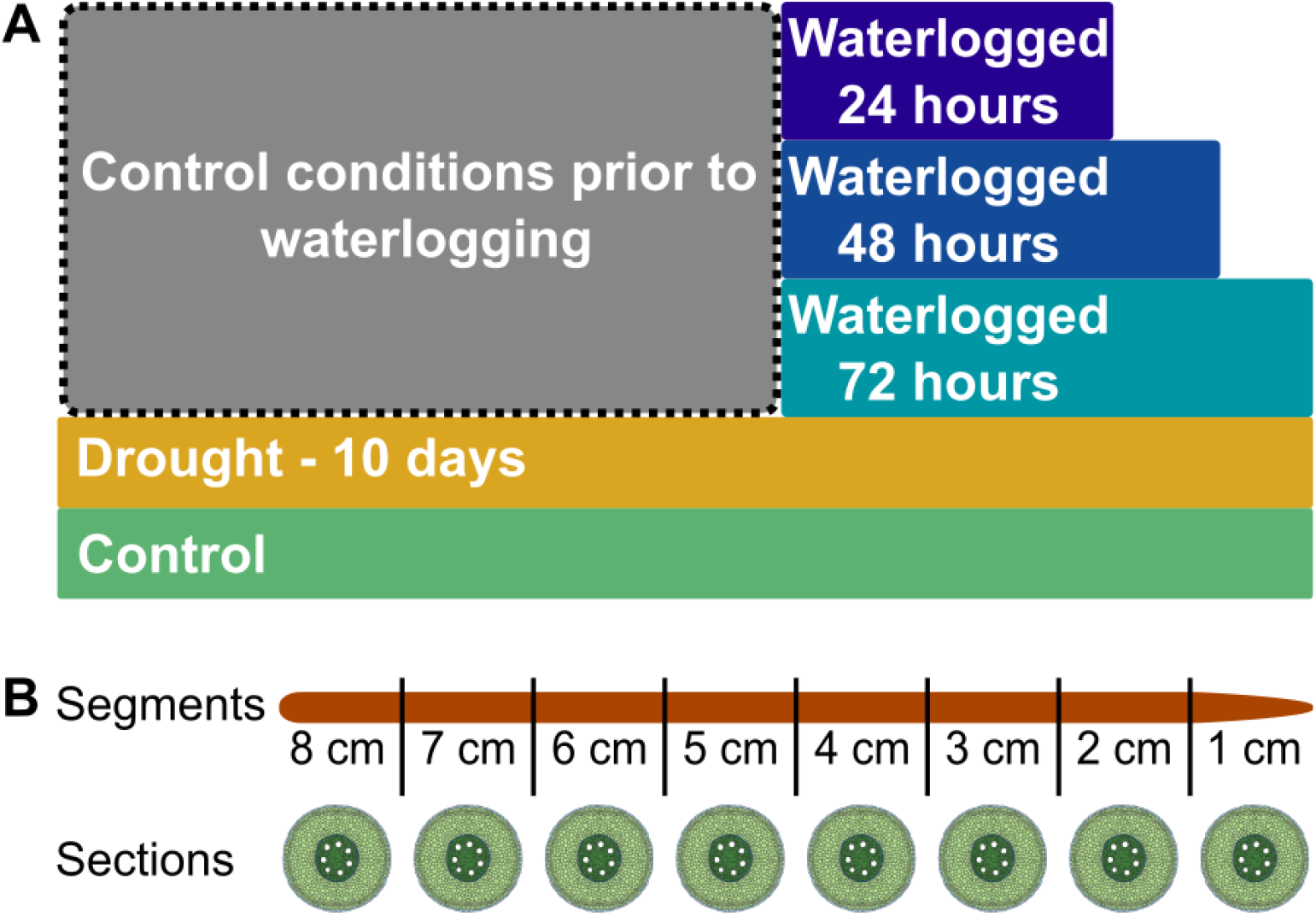
Experimental Design. Plants were grown for 30 days prior to splitting into **A)** treatment groups based on shoot height (n = 12 per treatment). **B)** A single nodal root was selected from each plant and segmented by centimeter prior to transverse sectioning, staining, and imaging.

### Root sectioning and image analysis

Root segments were hand sectioned transversally using a Teflon coated razor blade. Transverse sections were stained with 1.25% toluidine blue (MilliporeSigma, Burlington, MA, USA) for one minute, washed with 15% ethanol for 1 minute, and wet mounted on a slide with coverslip using distilled water. Sections were imaged at 40X magnification on an upright microscope (3000-LED) with an attached camera module (ExcelisHD/4K; ACCU-SCOPE Inc., Commack, NY, USA). For three plants per treatment, we made three sections per segment to assess the variation within the centimeter segment of root. We found that little variation was captured by collecting multiple sections per segment (see Results), thus we elected to collect only one section per segment for the remainder of the plants. Section images were analyzed using ImageJ (Schindelin *et al*., 2012); measurements collected are described in Table S1. All images were scaled to convert from pixel measurements to µm.

### Statistical Analysis

All statistical analysis was conducted in R v4.2.2 (R Core Team, 2022), utilizing the following packages tidyverse v2.0.0 (Wickham *et al*., 2019), lme4 v1.1-351 (Bates *et al*., 2015), emmeans v1.9.0 (Lenth *et al*., 2020), and ggpubr v0.6.0 (https://CRAN.R-project.org/package=ggpubr). To assess whether water stress treatments impacted the shoot system, we fit a linear mixed model both to shoot height and shoot dry weight, with treatment as a main effect and germination cohort as a random effect (accounting for transplant date). For each anatomical root trait, we fit the following linear mixed-effects model: *Root Trait ∼ Treatment + segment + Treatment × Segment + (1*|*Plant_ID)*. All models were assessed using a Type III ANOVA framework. Post hoc comparisons of estimated marginal means were made using a Dunnett’s test with each of the experimental treatments being compared to the control. For the 15 plants for which we measured three replicate sections within each centimeter segment, we also calculated the coefficient of variation for each root trait among replicate sections, root segments, and plants for each treatment to compare the importance of these sources of variation.

## Results

### The shoot was minimally impacted by short-term water stress

The average dry weight of the shoot 79.8 mg (SE = 4.74 mg) while the average height across treatments was 189.2 mm (SE = 6.5 mm). Water stress treatments did not significantly impact the height or dry weight of the shoot (Figure 2). This was expected, as the water stress treatments were short in duration, with no recovery period afterward, making a significant effect on shoot growth unlikely.

**Figure 2.**
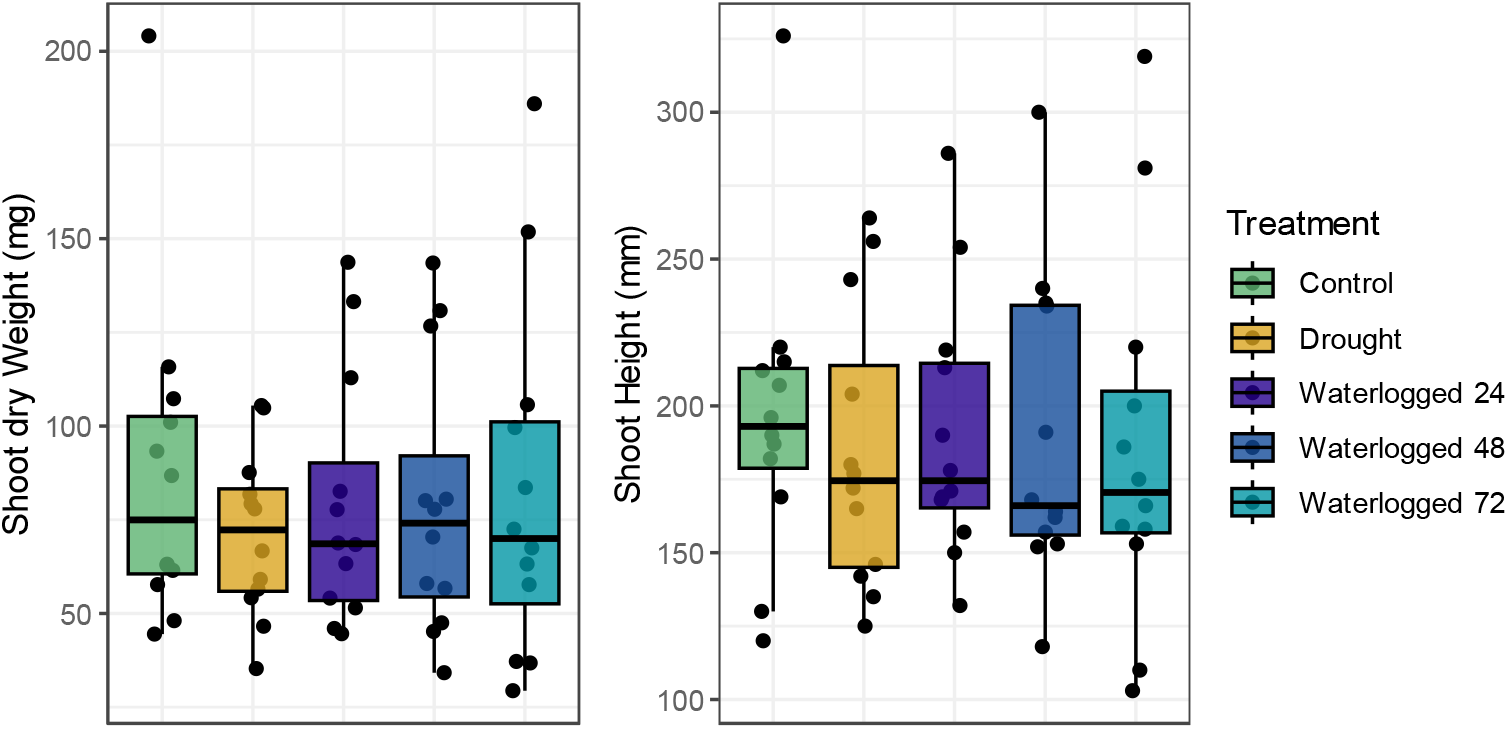
Shoot growth across water stress treatments. Box plots are colored by treatment for **A)** shoot dry weight (ANOVA; *‘Treatment’* p = 0.88, F_4,54_ = 0.11) and **B)** shoot height (ANOVA; *‘Treatment’* p = 0.98, F_4,54_ = 0.29). Boxplot hinges represent the 1^st^ and 3^rd^ quartiles; whiskers represent 1.5 times the interquartile range.

### Variation among plants and root segments dwarfs variation among technical replicates

To determine whether technical replicates are necessary to accurately quantify root anatomical traits, we quantified the variation between replicate sections made from a single centimeter segment. That is, for three plants per treatment, we performed sectioning in triplicate and compared the variation among replicate sections to the variation among segments (i.e., from root tip to base) and among plants within a treatment. Several traits stood out with high coefficients of variation, including root hair number and aerenchyma percentage (Figure 3). We observed that for all root traits, much more variation was captured among root segments and plants than among replicate sections (Figure 3B and C). This result motivated our decision to collect only a single section per segment of root for the remaining plants (N=44), which allowed us to increase the number of plants for each treatment group (N≥10 per treatment).

**Figure 3.**
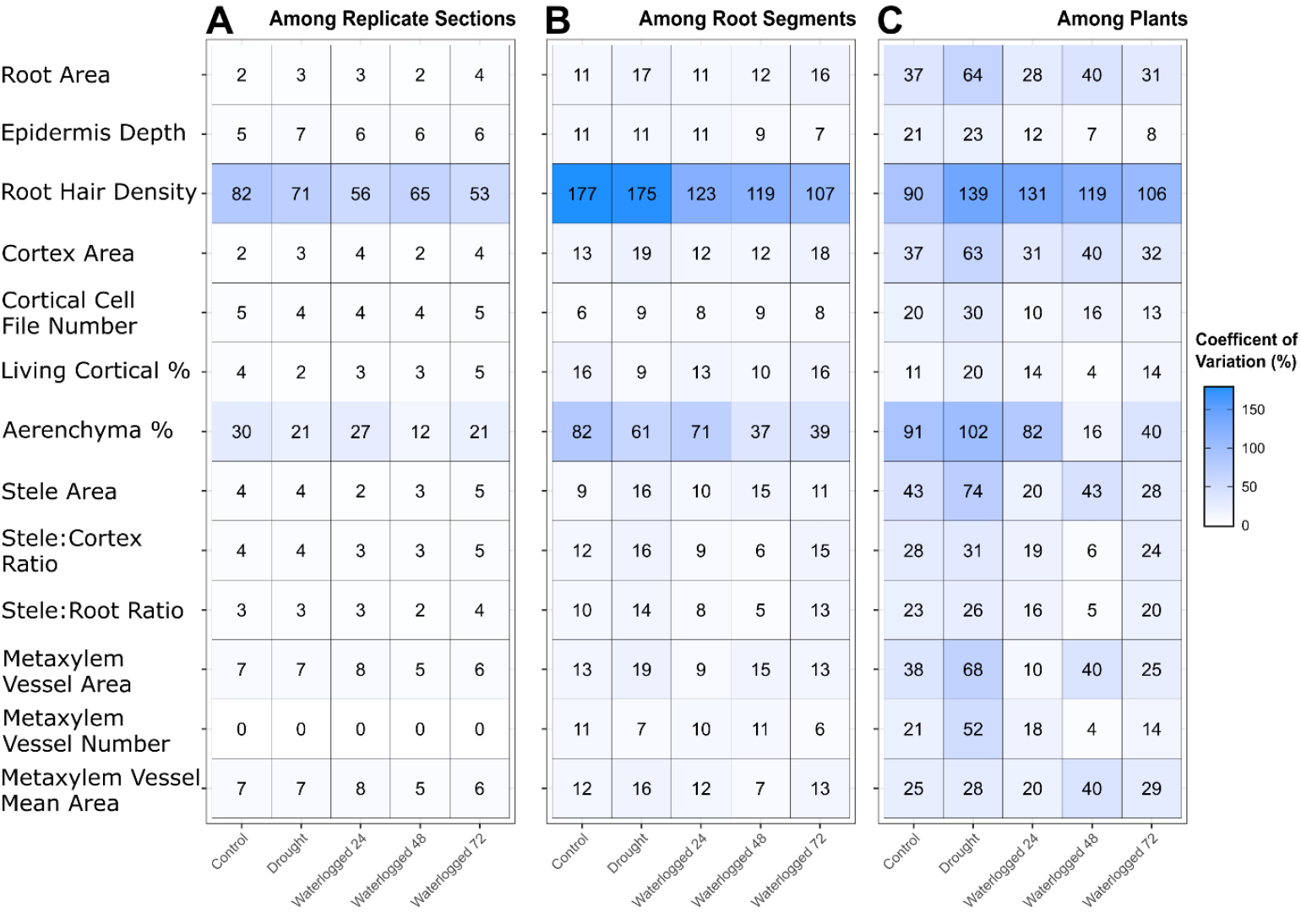
Little additional variation is captured by collecting replicate sections within a centimeter section of root. Coefficients of variation (CV) are displayed per trait and treatment combination **A)** among replicate sections, **B)** among root segments, and **C)** among plants. Shading is normalized across all panels to aid in comparison. Numbers in cells represent CV as percentages.

### Root anatomical responses depend on root segment and water stress treatment

Root trait responses differed according to which portion of the root was examined and the water stress treatment the plant experienced (Figure 4). Plant roots that had experienced drought showed a large increase in root hair number (Figure 4A), with an increase of 16.6 and 12.9 hairs/mm for the first two segments near the root tip, respectively. Waterlogged roots showed no significant deviation from the control plants in terms of root hair number (Figure 4B-D). In general, we observed that many trait responses were similar across all stresses; for example, the root area across segments was reduced under all stresses compared to the control, although this deviation was non-significant.

**Figure 4.**
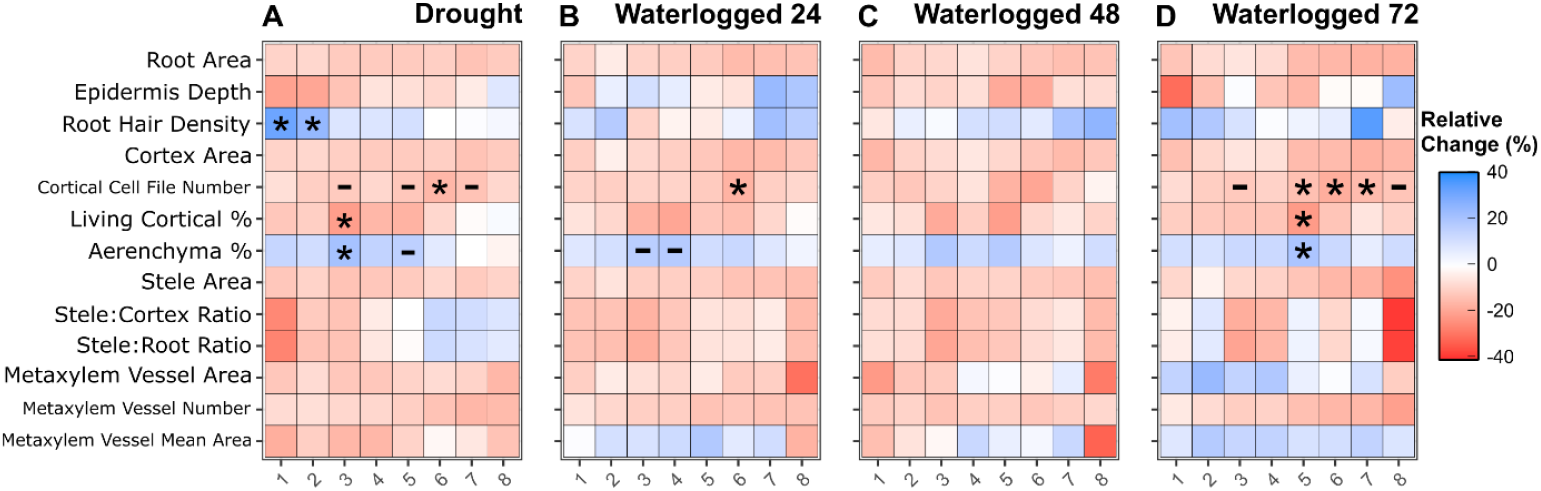
Water stress impacts root traits in differing ways. Heat maps depict the relative difference (%) in estimated marginal means between the control treatment and **A)** drought, **B)** waterlogged 24 hours, **C)** waterlogged 48 hours, and **D)** waterlogged 72 hours treatments. Significance was assessed using a Dunnett’s test (* *P-value*<0.05, - *P-value* <0.1).

For the root cortex we examined the cortical cell file number and the area of living cortex tissue. We found that under both drought and waterlogging, roots had a lower cortical cell file number across all root segments than control plants (13 vs 16, respectively). This was particularly evident 5-7 cm from the root tip, where both drought and waterlogging caused a significant reduction in cortical cell file number relative to the control (Figure 4 and 5A). In addition to fewer cortical cell files in the cortex, we observed the living cortical area (i.e., the area of the cortex made up of living cells) was decreased in our most extreme water stress treatments (Figure 5B). Both drought and 72 hours of waterlogging caused a significant reduction in living cortical area, but in different root segments (Figure 3A&D). Drought impacted the living cortical area most strongly 3 cm from the root tip while the 72-hour waterlogging showed the largest impact at 5 cm from the root tip. In both cases the decreased living cortical area was due to an increase in aerenchyma tissue in the cortex.

**Figure 5.**
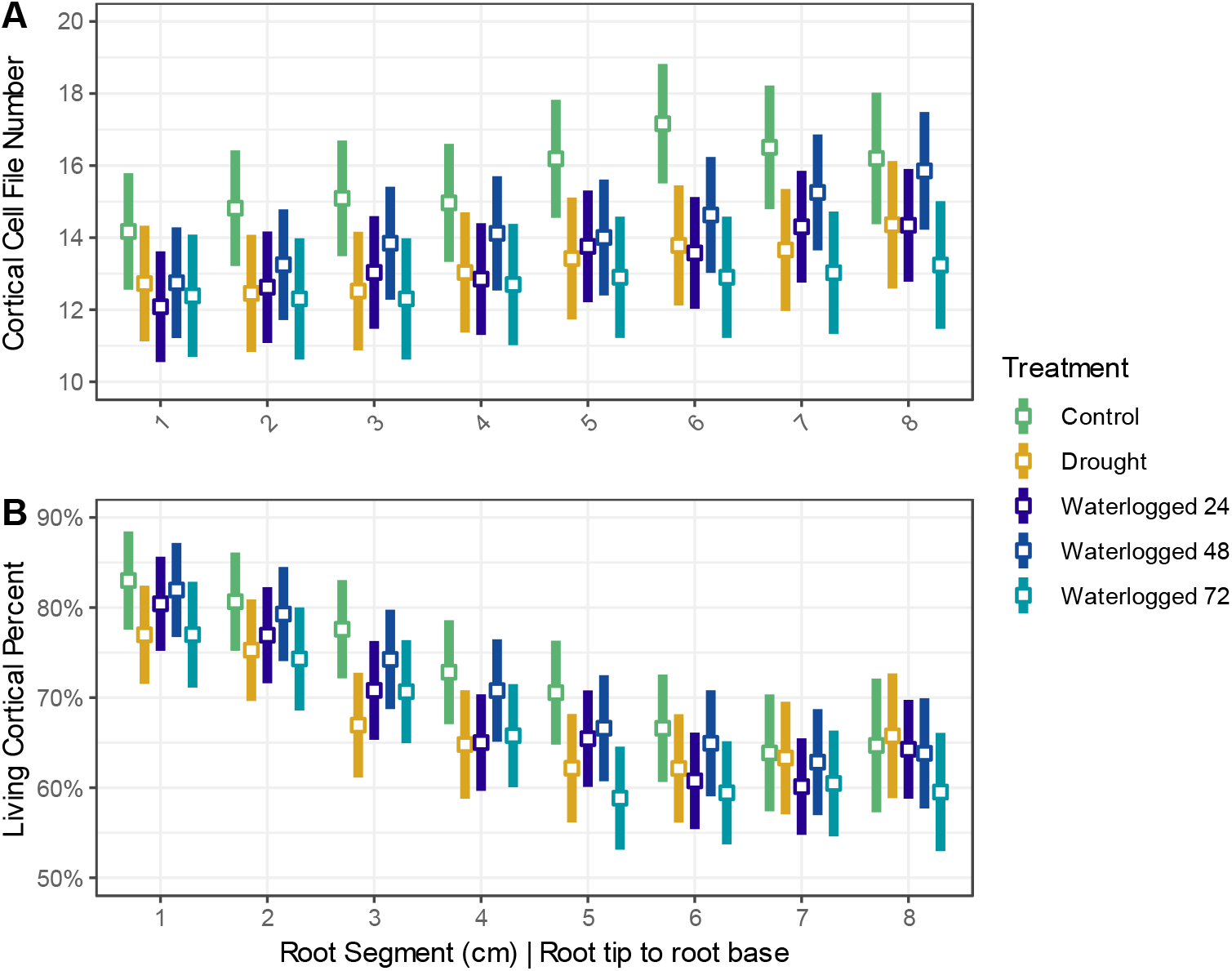
The root cortex is reduced in cortical cell layers and living cells under stress. Line plots depict the estimated marginal means (white squares) with 95% upper and lower confidence intervals for **A)** cortical cell file number and **B)** the proportion of the cortex composed of living cells.

### Drought and waterlogging have contrasting effects on stele anatomy

The stele contains the vascular tissue of the root and represents an important conduit for both water and solute transport between roots and shoots. Similar to our observations for total root area, the stele area of stressed plants was on average smaller than the control (figure 6A). The number of metaxylem vessels was also reduced in stressed plants by ∼1 to 1.5 per plant. Plants that were waterlogged for 72 hours exhibited the largest vessel areas (1932 μm^2^, SE = 187), suggesting a strategy of producing larger but fewer in number metaxylem vessels (Figure 6B&C). This contrasted with drought-stressed plants which had metaxylem vessel counts similar to those waterlogged for 72 hours but exhibited the smallest vessel areas (1373 μm^2^, SE = 181).

**Figure 6.**
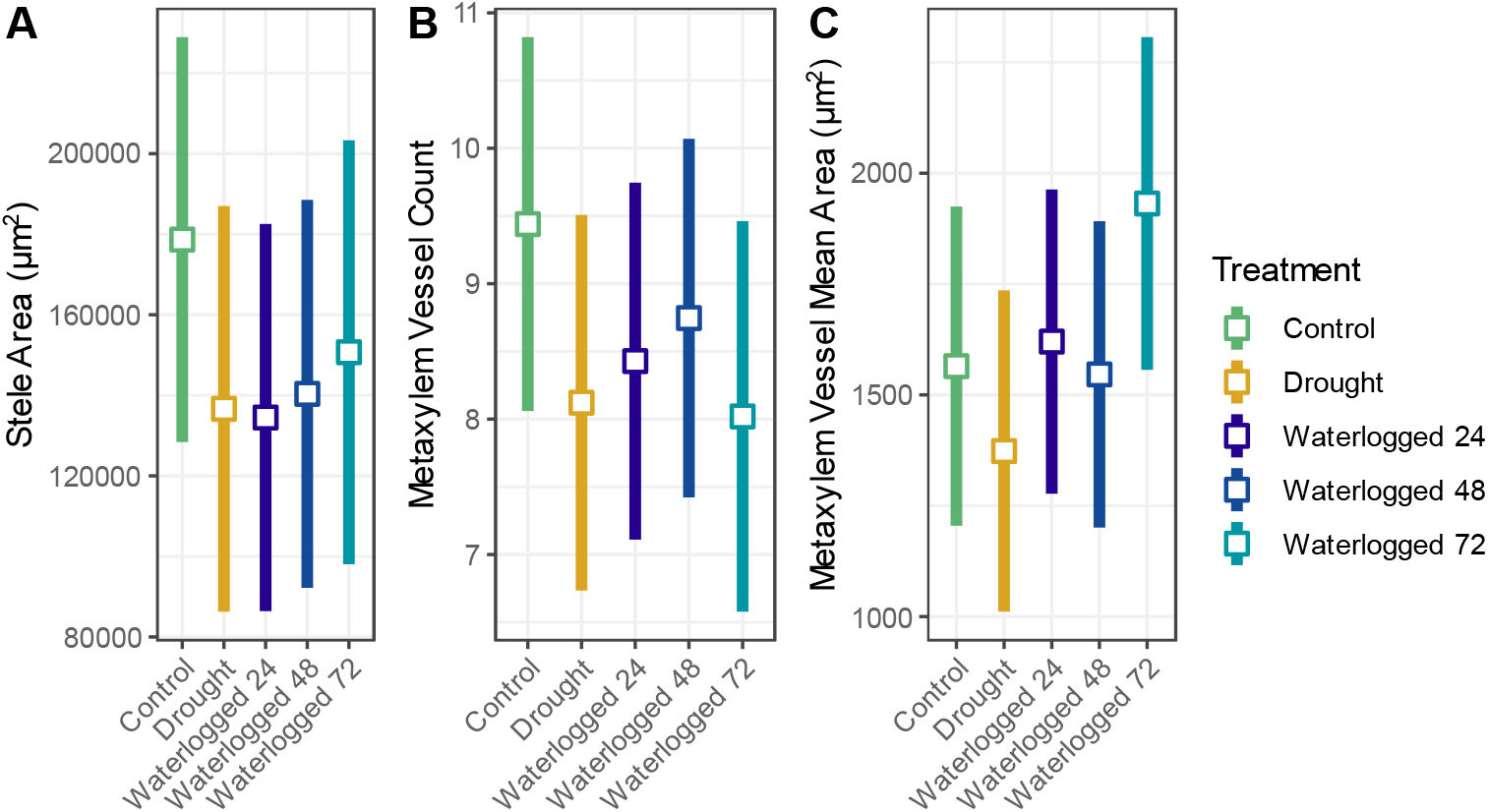
Metaxylem anatomical responses to water stress. Line plots depict the estimated marginal means (white squares) with 95% upper and lower confidence intervals for **A)** stele area, **B)** metaxylem vessel count, and **C)** metaxylem vessel mean area.

We observed a pattern in the ratio between the stele and root area where, at either end of the water stress extreme (drought vs 72-hour waterlogging), the portion of the root composing the stele was either reduced or enlarged (Figure 4A&D). To examine our two most extreme treatments (drought and 72-hour waterlogging) in greater detail, we removed the less intense waterlogging durations. We found a significant interactive effect (p = 0.02, F_14,157_ = 2.01), with the water stress treatments impacting different portions of the root (Figure 7). At the tip of the root, droughted plants showed a reduced stele size in relation to the root size (14.5% of the root area vs 16.5% and 16.3% for control and waterlogging, respectively) while the control and waterlogged treatments were equivalent (Figure 7). The opposite was the case for the segment furthest from the root tip, where the waterlogged treatment showed a reduction in stele size (1.8-2.3% reduction from control and drought treatments, respectively).

**Figure 7.**
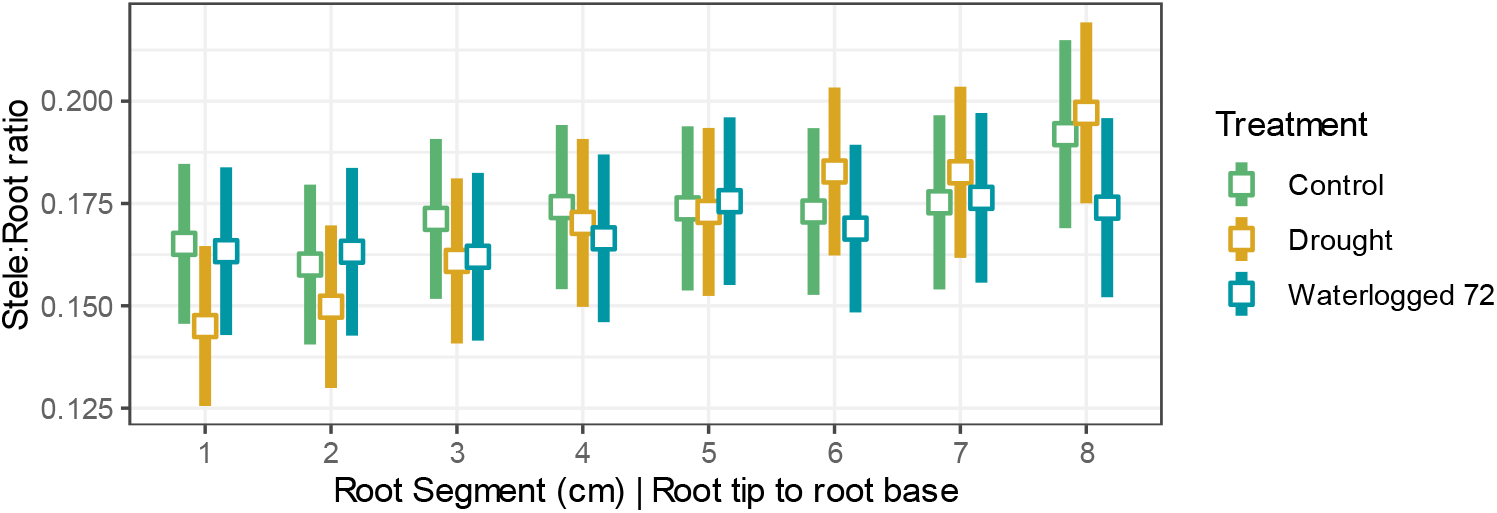
The root stele shows contrasting patterns to water stress across the root length. Line plots depict the estimated marginal means (white squares) with 95% upper and lower confidence intervals for the Stele:Root Ratio.

## Discussion

Despite all that is known about root anatomical responses to stress, a lack of resolution along the root for where different phenotypic responses are the strongest hinders integration with other data types where sampling effort is limiting (e.g., microbiome, transcriptome, proteome). Integrative approaches such as these typically utilize tens to hundreds of plants per experiment, creating a considerable barrier as root anatomical measurements require considerable time to acquire. Here, we detail for *Tripsacum dactyloides* the traits and locations along the length of the root that vary as a response to water stress, providing the necessary information to explore more targeted experiments.

Commonalities exist in root anatomical traits at water stress extremes. While reduced cortical cell file number have been observed previously for Poaceae under drought stress (Fraser *et al*., 1990; Chimungu *et al*., 2014a; Colombi *et al*., 2019), we observed this same phenomenon not only in droughted roots but also in waterlogged roots (Fig. 5A). The reduction in cortical cell file number occurred within a common portion of the root, around 5-7 cm from the root tip, and was evident even in roots that experienced just 24 hours of waterlogging. This indicates a more general stress response, whereby plants suppress root enlargement by reducing cortical cell file addition under water stress. A reduced cortical cell file number and increased cortical cell diameter have been correlated with reduced energy costs for roots in wheat (Colombi *et al*., 2019). Maize genotypes with reduced cortical cell file numbers and larger cortical cell sizes also exhibited improved yields under water limitation in field settings (Chimungu *et al*., 2014a,b). Although cell size was not quantified in this study, cortical cell files in the middle of the cortex, especially those flanked by aerenchyma lacunae, appeared qualitatively larger in size for stressed plants. Cortical aerenchyma formation also showed a common response between water stress extremes in *T. dactyloides*, with significant increases in the aerenchyma percentage of the root cortex 3-5 cm from the root tip under both drought and waterlogging (Fig. 4A). Cortical aerenchyma formation decreases the energy costs for sustained root growth, which under drought can allow for greater soil exploration (Zhu *et al*., 2010; Jaramillo *et al*., 2013) and under waterlogging can serve as an internal gas space allowing oxygen diffusion between the shoot and root system (Jackson & Armstrong, 1999; Colmer, 2003; Abiko & Miyasaka, 2020; Pedersen *et al*., 2021a). The amount of aerenchyma formed for *T. dactyloides* under flooding is similar to nodal roots of *Zea nicaraguensis*, a wild relative of maize, which was previously observed at >20% aerenchyma (as a fraction of the whole root area) for root segments 6 cm from the tip (Gong *et al*., 2019). Of note, for *Z. nicaraguensis* aerenchyma formation was not apparent within the first 1-2 cm from the root tip with 72 hours of stagnant conditions (Gong *et al*., 2019), whereas for *T. dactyloides*, we observed aerenchyma formation closer to the root tip, as close as 1 cm from the tip. Both species produced aerenchyma under control conditions, indicating constitutive production which is linked to improved performance under drought and waterlogging conditions, as has been observed for both species (Ray *et al*., 1998; Clark *et al*., 1998; Mano & Omori, 2013).

Other traits, however, showed contrasting responses to the water stress extremes. For instance, droughted plants—but not waterlogged plants—had significantly more root hairs in the first 1-2 cm of the root. This is in line with previous research on drought stress in plants (Mackay & Barber, 1985; Carminati *et al*., 2017; Marin *et al*., 2021), with a hypothesized role for root hairs in water uptake, although this is dependent on both soil texture and species (Cai *et al*., 2021; Duddek *et al*., 2022). In contrast, root hair number of waterlogged plants was similar to that of control plants. We note that although we were able to count root hairs in most cross-sections, some images presented difficulty due to lack of focus at the periphery of the root endodermis and occlusion in samples with copious root hairs, both of which made exact enumeration challenging. In these cases, we enumerated as well as possible. Droughted plants tended to have the most occluded root hairs due to high densities in certain segments. It would be preferable in future experiments to measure root hair density by imaging the root section longitudinally, either prior to cross sectioning or on another suitable root, to allow for a greater area to be measured. Given that our results match what has been observed in other species, however, we believe this methodological issue does not invalidate our data.

Metaxylem vessels also exhibited contrasting patterns under water stress. While the number of metaxylem vessels was reduced to a similar extent across treatments, droughted roots showed smaller vessel areas in comparison to the most extreme waterlogging treatment. Plasticity in metaxylem vessel diameter is associated with increased performance under drought stress as narrower vessels are better able to conduct water under as soils dry and water potential becomes increasingly negative (Richards & Passioura, 1989; Vasellati *et al*., 2001; Kadam *et al*., 2015; Prince *et al*., 2017). In monocots, metaxylem of the seminal roots with smaller diameters have been found to be less susceptible to cavitation (Klein *et al*., 2020; Harrison Day *et al*., 2023). Compared to drought stress, less is known about the response of metaxylem vessels to waterlogging. We found that roots that experienced the longest duration of waterlogging had the largest metaxylem vessels, but this contrasts with other studies that have found an opposite pattern or no effect of waterlogging on metaxylem vessel size in other species (Huang *et al*., 1994; Vasellati *et al*., 2001).

Another limitation to our experiment is the use of only a single cultivar, ‘Pete’, which may limit the transferability of the results to other accessions and related species. ‘Pete’ is expected to be highly heterozygous as it was generated from a composite of 70 wild-collected accessions and has been maintained as an open-pollinated population, so it may exhibit higher phenotypic variance than would be observed for a clonal wild-collected accession. Little research has focused on root anatomical trait variation at the intra- and inter-specific level in *Tripsacum* (Ray *et al*., 1998) compared to its relatives in the genus *Zea* (Burton *et al*., 2013; McLaughlin *et al*., 2024). As the closest temperate relative of *Z. mays*, with which it can interbreed, (Berthaud *et al*., 1997; Ray *et al*., 1999; Yan *et al*., 2020), the exploration of a greater diversity of *T. dactyloides* genotypes could be beneficial to breeding programs. For instance, root anatomical traits are known to mediate nutrient acquisition and stress tolerance (Kadam *et al*., 2015; Klein *et al*., 2020; Lynch *et al*., 2021) and the introgression of *T. dactyloides* loci into the genomes of *Z. mays* and *Z. nicaraguensis* has improved hypoxia tolerance in those species (Mano *et al*., 2007, Mano *et al*., 2009; Gong *et al*., 2019). Furthermore, genetic variation within *Tripsacum* reflects adaptation to local environmental conditions and can suggest targets for genetic improvement within the maize genome, as has been shown for cold tolerance genes that evolved in parallel in the two species (Yan *et al*., 2019). Therefore, understanding genetic variation for root anatomy among *T. dactyloides* accessions from diverse habitats is a promising avenue of future research that could accelerate the development of more stress-resistant maize varieties.

Roots, both externally and internally, serve as diverse habitats for microorganisms. Stress-induced changes in root anatomy, even if small in magnitude, can have large implications for interactions that take place at the microscopic level between plants and microorganisms. Associations formed between plants and microorganisms can have profound effects on plant responses to abiotic stressors (Xu *et al*., 2018a; Durán *et al*., 2018; Rolfe *et al*., 2019), with some microorganisms able to influence key plant stress hormones that underlie rapid anatomical responses (Ravanbakhsh *et al*., 2017, Ravanbakhsh *et al*., 2018; Xu *et al*., 2018b). To the extent that root anatomical traits or their responses to stress are genetically controlled, they may represent one mechanism explaining the genetic variation for colonization rates of root-associated fungi that has been observed in *T. dactyloides*. Fungal colonization rates were negatively correlated with mean annual precipitation at the *T. dactyloides* accessions’ original collection sites (Kural-Rendon *et al*., 2023), consistent with a genetic link between adaptation to water stress and interactions with fungi. It is interesting to postulate that stress-induced changes in root anatomy may also modulate the conditions experienced by microorganisms under abiotic stress. The relationship between root anatomy and microbial interactions is an emerging area of research interest (Birt *et al*., 2022; Kawa & Brady, 2022; Galindo-Castañeda *et al*., 2024). For example, in a panel of maize genotypes living cortical area was negatively associated with colonization by arbuscular mycorrhizal fungi, but positively associated with pathogenic fungi (Galindo-Castañeda *et al*., 2019). Bacterial community composition is also partially explained by the amount of root cortical aerenchyma in maize (Galindo-Castañeda *et al*., 2023). Both between root types (seminal and nodal roots) and longitudinally along a single root, bacterial and fungal communities show remarkable compositional variation (Rüger *et al*., 2021), similar to the degree of variation observed between different plant species (Kawasaki *et al*., 2021). Root tips are particularly diverse in microorganisms, with diversity levels for the root rhizoplane/endosphere similar to that of bulk soil (Kawasaki *et al*., 2021). Microbial niches also change across the soil-root continuum, with large compositional changes from the rhizosphere to the endosphere (Edwards *et al*., 2015), and to a smaller degree changes between periderm, phloem and xylem tissue (Zhou *et al*., 2022). By using information on which root regions exhibit the greatest stress-mediated responses, targeted techniques such as metatranscriptomics can be utilized to assess the response of microorganisms to water stress at a scale that has rarely been explored. This work also facilitates reductionist lines of inquiry into the impact of individual and synthetically assembled microbial communities on plant root traits and vice versa.

## Code availability

All code to reproduce the analysis and figures is available on GitHub at https://github.com/Kenizzer/Tripsacum_dactyloides_Root_Anatomical_Responses_to_Drought_and_Waterlogging. Light microscopy images of *Tripsacum dactyloides* roots are available at https://doi.org/10.5281/zenodo.14834918.

## Acknowledgements

We would like to thank Natalie Ford and Martel Ellis for assistance planting and caring for *Tripsacum*. This work was funded by the National Science Foundation. DT was funded as part of a research experience for undergraduates’ internship made possible by an NSF Biology Integration Institute grant #2120153 and JFS was funded by a postdoctoral research fellowship #2305703. MRW was supported by NSF grant IOS-2016351.

## Author Contributions

JFS and MRW designed the experiment. JFS performed experimental treatments and root harvesting. JFS, DT, LB, SA, performed root sectioning and image analysis. JFS analyzed data. JFS wrote the first draft of the manuscript which all authors read, commented on, and edited.

## Supplemental Figures

**Figure S1.**
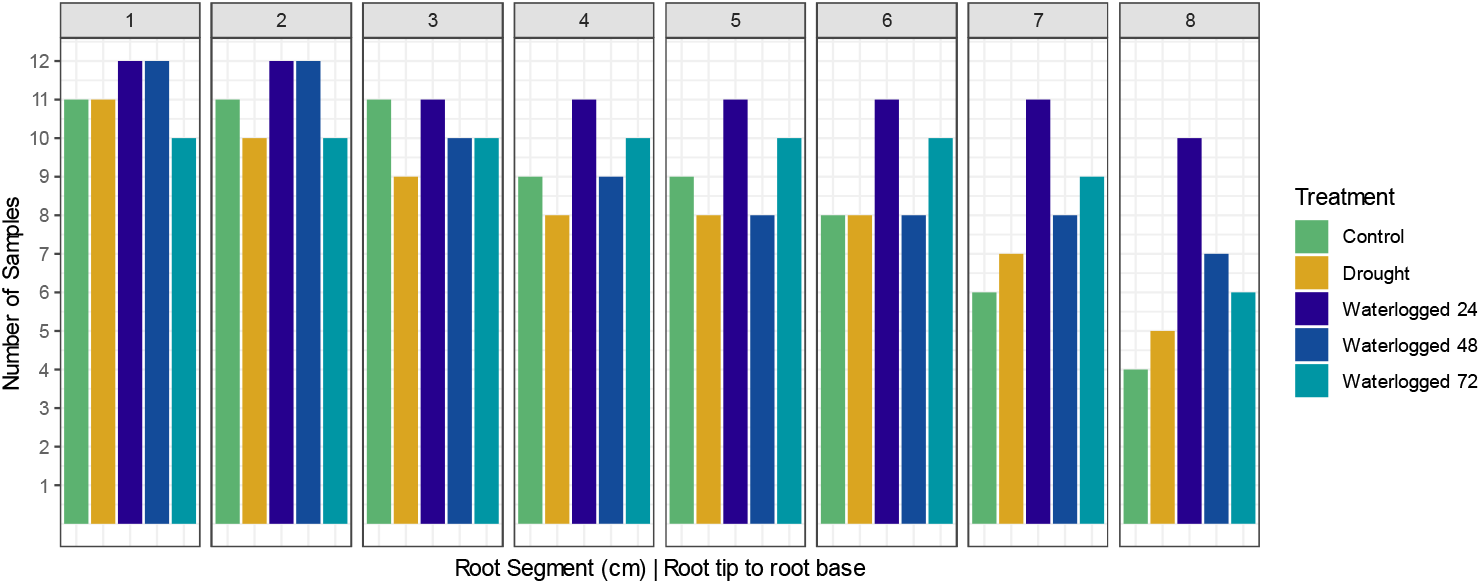
Root segment counts by treatment. Each sample was divided into up to eight segments (facet panels), some plants had roots shorter than eight centimeters (n = 24) while several plants did not have nodal roots suitable for collection (n = 4).

## Supplemental Tables

**Table S1:**
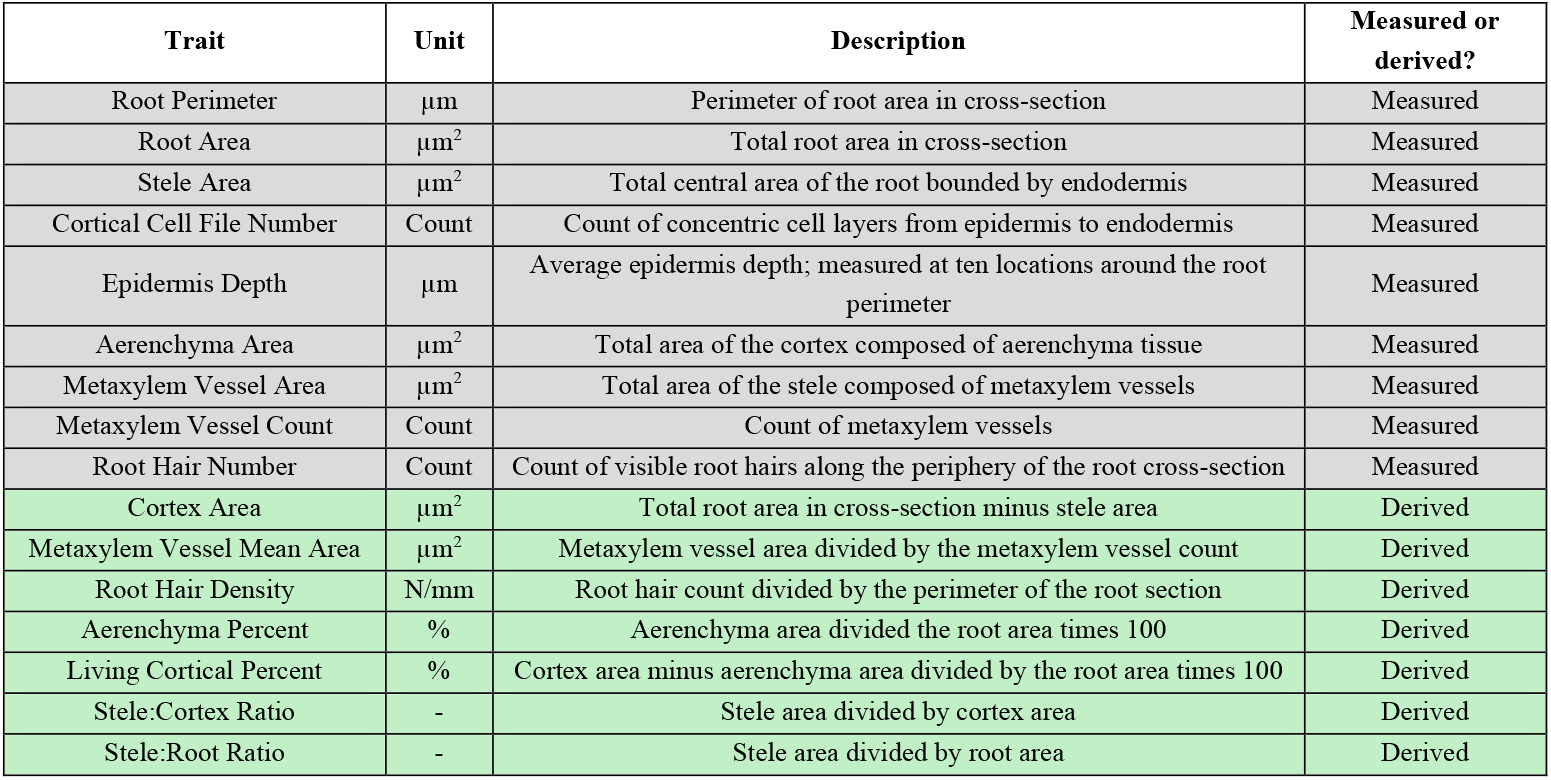
Root trait measurements collected from root cross-sections.

| Trait | Unit | Description | Measured or derived? |
| --- | --- | --- | --- |
| Root Perimeter | $\mu\text{m}$ | Perimeter of root area in cross-section | Measured |
| Root Area | $\mu\text{m}^2$ | Total root area in cross-section | Measured |
| Stele Area | $\mu\text{m}^2$ | Total central area of the root bounded by endodermis | Measured |
| Cortical Cell File Number | Count | Count of concentric cell layers from epidermis to endodermis | Measured |
| Epidermis Depth | $\mu\text{m}$ | Average epidermis depth; measured at ten locations around the root perimeter | Measured |
| Aerenchyma Area | $\mu\text{m}^2$ | Total area of the cortex composed of aerenchyma tissue | Measured |
| Metaxylem Vessel Area | $\mu\text{m}^2$ | Total area of the stele composed of metaxylem vessels | Measured |
| Metaxylem Vessel Count | Count | Count of metaxylem vessels | Measured |
| Root Hair Number | Count | Count of visible root hairs along the periphery of the root cross-section | Measured |
| Cortex Area | $\mu\text{m}^2$ | Total root area in cross-section minus stele area | Derived |
| Metaxylem Vessel Mean Area | $\mu\text{m}^2$ | Metaxylem vessel area divided by the metaxylem vessel count | Derived |
| Root Hair Density | N/mm | Root hair count divided by the perimeter of the root section | Derived |
| Aerenchyma Percent | % | Aerenchyma area divided the root area times 100 | Derived |
| Living Cortical Percent | % | Cortex area minus aerenchyma area divided by the root area times 100 | Derived |
| Stele:Cortex Ratio | - | Stele area divided by cortex area | Derived |
| Stele:Root Ratio | - | Stele area divided by root area | Derived |

